# MNRR1 is required for Triple Negative Breast Cancer growth and metastasis and can be targeted by repurposed drugs

**DOI:** 10.64898/2026.01.31.703055

**Authors:** Mrunal Sathe, Paige Minchella, Caleb Vegh, Mithra Modikuppam Dharmalingam, Vignesh Pasupathi, Yue Xi, Sagarika Rai, Mrunmayee Patil, Habiba Elshenawy, Sydney Rudolph, Gaurav Bhatti, Adi L. Tarca, Neeraja Purandare, Andrew Fribley, Lawrence I. Grossman, Siddhesh Aras

## Abstract

Despite progress in recent decades, breast cancers remain the most diagnosed malignancies, and second leading cause of cancer death, in women. Although improved screening and systemic and endocrine adjuvant approaches have contributed to major declines in breast cancer mortality, current standard of care drugs are extremely toxic, and many women continue to be overtreated. Although nearly two-thirds of breast cancers are hormone-responsive, aggressive subtypes, particularly Triple Negative Breast Cancer (TNBC), still lack safe oral medications. Recently the roles that mitochondria play in TNBC carcinogenesis, metastasis, and resistance to treatment have garnered a great deal of attention. Contrary to the popular dogma that cancer cells are powered by glycolysis, metastatic breast cancer cells have enhanced mitochondrial function. Our work identified that Mitochondrial Nuclear Retrograde Regulator 1 (MNRR1; also, CHCHD2, PARK22), a key coordinator of mitochondrial-nuclear crosstalk that is physically present in both compartments, is overexpressed in TNBC cells and is an important regulator of metastasis signaling. We have identified Heat Shock Factor 1 (HSF1) as the main transcription factor that activates the *MNRR1* promoter in TNBC cell lines. In the mitochondria, MNRR1 protein facilitates ATP production and inhibits apoptosis, whereas in the nucleus it regulates the transcription of stress-responsive genes including several required for epithelial to mesenchymal transition (EMT), metabolic flexibility, and cell growth. Thus, each of the bi-organellar functions of MNRR1 constitutes processes regarded as hallmarks of cancer. For reasons that are not yet fully understood, MNRR1 levels display a significant and robust ancestry bias, showing increased expression in tumor samples from Non-Hispanic Black (NHB) women when compared to disease-matched tumors from Non-Hispanic White (NHW) patients. It is possible that increased levels of MNRR1 may underlie the aggressive metastatic phenotype observed in many NHB patients. In further support of this observation, loss of MNRR1 function, either genetically or by use of inhibitors, reduces TNBC growth and metastasis. MNRR1 therefore is an attractive therapeutic target that could be exploited for design of novel therapies or as adjuncts to existing ones.

## Introduction

Although the rates of breast cancers appear to have stabilized in many developed countries over the past few decades, developing counties have not realized this benefit, and it is still the second leading cause of cancer-related deaths in women around the globe [1]. According to the latest statistics, United States is still anticipating 324,580 new breast cancer cases with more than 42,000 deaths in 2026 [2]. While chemotherapeutic regimens have reduced the previous mortality rates, toxicity, resistance, and often debilitating late effects continue to provide unacceptable morbidity and mortality [3]. Triple Negative Breast Cancer (TNBC), shouldering a small proportion of the disease burden, runs an aggressive course associated with higher mortality rates [4]. Tumor heterogeneity and lack of specific targets have contributed to the use of cytotoxic drugs as the frontline therapy with only about 50% of patients achieving a complete response [5]. Therefore, there is an urgent unmet need for the identification of specific targeted therapies or adjuvants that would be both effective and less toxic. Additionally, TNBC exhibits an ethnic bias with Non-Hispanic Black (NHB) women incurring higher rates with a significantly higher mortality risk compared to Non-Hispanic White women (NHW) [6, 7].

Mitochondria, the cellular hubs for energy production, generate reactive oxygen species (ROS) as signaling molecules and have been shown to play a vital role in myriad aspects of cancer cell metabolism. The importance of mitochondria in tumor cells was uncovered in seminal studies showing defective tumor growth upon deletion of mitochondrial DNA (mtDNA), or the mitochondrial transcription factor (TFAM) [8, 9]. Similarly, highly metastatic cells display an increase in mitochondrial biogenesis and respiration [10]. This enhanced mitochondrial function was shown to provide energy to the migrating cells. Finally, multiple studies have revealed that mitochondria play key roles in cancer cell pathways that favor tumorigenesis such as impaired apoptosis, metabolic substrate flexibility, and ROS signaling (reviewed in [11]). The pivotal roles mitochondria assume in tumorigenesis and death make them attractive targets for cancer therapeutics. Mitochondrial Nuclear Retrograde Regulator 1 is a highly conserved bi-organellar protein localized to both mitochondria and the nucleus [12]. In mitochondria, phosphorylated MNRR1 can interact with cytochrome *c* oxidase (COX) to activate oxidative phosphorylation (OxPhos) or interact with Bcl-xL to prevent apoptosis. In the nucleus, MNRR1 can be deacetylated by Sirt1 to promote its functioning as a transcription regulator. Loss-of-function mutations in MNRR1 are associated with a neurodegenerative phenotype [13]. Cells deficient for MNRR1 display reduced growth, enhanced ROS, and a fragmented mitochondrial phenotype [12, 14]. Transcript levels of *MNRR1* are associated with a cancerous phenotype [15]. We have previously shown MNRR1 levels to be higher in multiple breast cancer cell lines used in research [14]. Furthermore, in ovarian cancer, MNRR1 was shown to be sufficient to induce spheroid formation [16]. Similarly, MNRR1 supported tumor growth and metastasis of glioblastoma [17], renal cell carcinoma [18], and hepatocellular carcinoma [19]. Finally, increased levels of MNRR1 have been associated with lung adenocarcinomas with an unfavorable outcome [20].

The current study has been focused on a more detailed characterization of the mechanisms in each organelle by which MNRR1 regulates growth and metastasis of TNBC using *in vitro* and *in vivo* models. Mitochondrial MNRR1 activates OxPhos to support the bioenergetic needs of proliferating cells while further driving the malignant phenotype by inhibiting apoptosis. Nuclear MNRR1 induces the Epithelial-Mesenchymal Transition (EMT) program to facilitate invasion and metastasis. We also report that high MNRR1 levels may underlie the ethnic bias observed in TNBCs. Finally, we show that small molecule inhibitors of MNRR1 identified on a drug screen can be repurposed as therapeutics or adjuvants in the treatment of TNBC.

## Results

### MNRR1 levels are higher in Triple Negative Breast Cancer cells and support cancer cell growth

TNBC cells contain higher levels of MNRR1 than control cells. The increased protein levels could result from increased transcription or enhanced post-translational protein stability. To parse this with respect to MNRR1, we used an isogenic pair of TNBC cell lines: HTB125 (non-malignant) and HTB126 (TNBC). *MNRR1*-promoter luciferase levels in the HTB126 cancer cell line displayed an ∼15-fold induction compared to the non-cancerous control HTB125 (**Fig. 1A**). Furthermore, the increase in *MNRR1* transcription led to an increase at the protein level (**Fig. 1A**). As we previously showed, increased MNRR1 levels are associated with enhanced mitochondrial function [12, 14, 21, 22]. HTB126 cells displayed a significant increase in mitochondrial oxygen consumption rate (OCR) compared to the HTB125 cells (**Fig. 1B**). We have previously shown that *MNRR1* transcript levels are increased in a panel of breast cancer cell lines that included TNBCs [14]. Similar to what was observed with the isogenic pair, increased MNRR1 transcript and protein levels were observed in two additional TNBC cell lines, MDA-MB-231 (231) and MDA-MB-468 (468), when compared to MCF10A (10A) controls (**Fig. 1C and 1D**). Although both the TNBC cell lines displayed increased MNRR1 compared to 10A, a 3-4-fold variation was observed between 231 and 468 cells (**Fig. 1D**), highlighting that there is variability even between TNBC patients. In addition to transcript and protein levels of MNRR1, the same TNBC cell lines also displayed a parallel increase in respiratory function (**Fig. 1E**). The enhanced oxidative phenotype has also been previously reported in TNBC cell lines [23]. In the current study, increased MNRR1 protein was also observed in TNBC cells measured by immunofluorescence microscopy (**Supplementary Fig. S1a and S1b**). As MNRR1 supports cell growth [12], and cancer by definition is an uncontrolled growth of cells, we assessed the ability of MNRR1 to increase the 10A (non-malignant breast) growth rate. 10A cells overexpressing MNRR1 (WT-R1) have significantly higher growth compared to those expressing an empty vector (EV) when measured after 48 or 72 hrs post-transfection (**Fig. 2A**).

**Figure 1:**
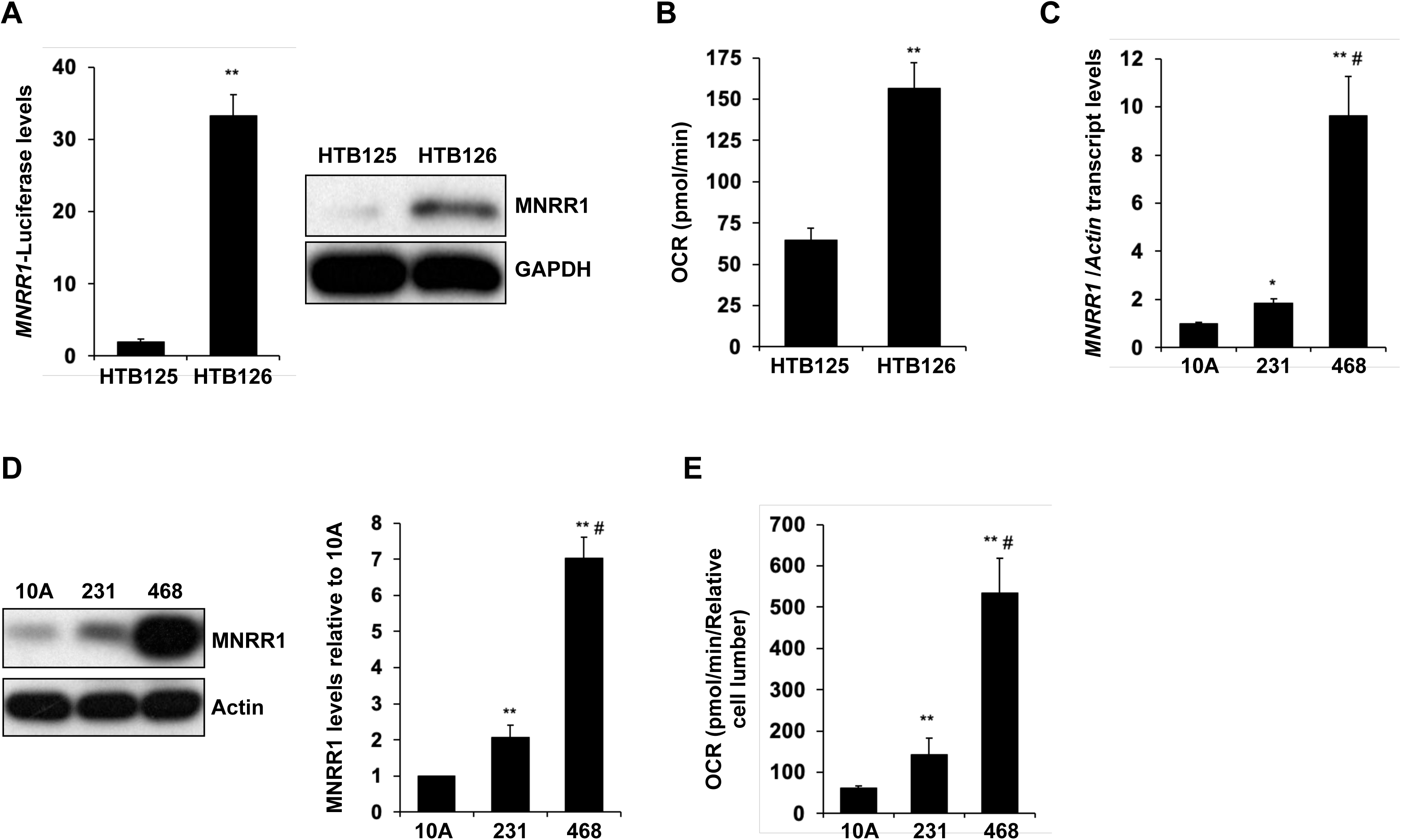
MNRR1 levels are higher in Triple Negative Breast Cancer cells. **A.** *Left,* Relative activation of *MNRR1*-promoter harboring luciferase reporter is higher in the breast cancer line HTB126 cells as compared to the isogenic control line HTB125 cells (n=6, **p<0.01). *Right,* MNRR1 protein levels are higher in cells as compared to HTB125 cells analyzed using western blot. GAPDH was probed as loading control. **B.** Basal mitochondrial oxygen consumption in cells is higher than HTB125 cells (n=10, **<0.01). **C.** Transcript levels of endogenous *MNRR1* are higher in 231 and 468 cells as compared to 10A cells (n=6, *p<0.05 for 231, **p<0.01 for 468 compared to 10A, **^#^**p<0.01 comparing 231 vs 468). *Actin* was used as housekeeping gene in RT-qPCR analysis. **D.** *Left,* MNRR1 protein levels are higher in in 231 and 468 cells as compared to 10A cells analyzed using western blot. Actin was probed as loading control. *Right,* quantitation of MNRR1 protein levels in MCF10A, 231, and 468 cells (n=5, **p<0.01 for 231 and 468 compared to 10A, **_#_**p<0.01 comparing 231 vs 468). **E.** Basal mitochondrial oxygen consumption in cells is higher than MCF10A (n=at least 8, **p<0.01 for 231 and 468 compared to 10A, **^#^**p<0.01 comparing 231 vs 468).

**Figure 2:**
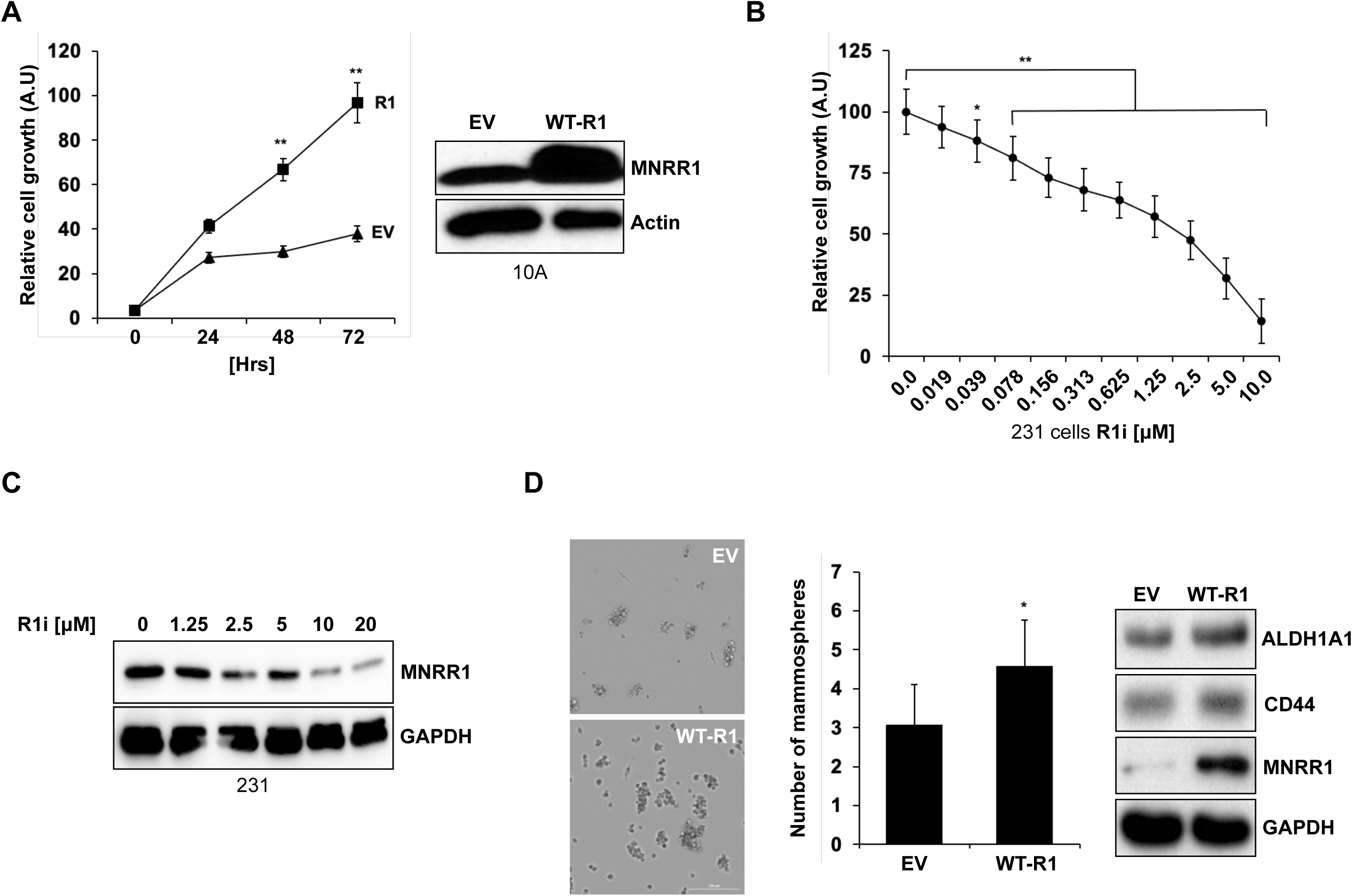
Higher MNRR1 levels support enhanced cell growth in breast cancer cells. **A.** *Left,* Cell growth of 10A overexpressing MNRR1 is higher that empty vector (EV) overexpressing cells measured using RealTime-Glo™ luminescence assay over 72 hrs (n=5, **p<0.01). *Right,* Western blot to confirm overexpression of MNRR1 in MCF10A cells. Actin was probed as loading control. **B.** Cell growth is inversely proportional to increasing concentrations of MNRR1 inhibitor (R1i, lovastatin) in 231 cells treated for 36 h (n= 8, *p<0.05, **p<0.001). **C.** MNRR1 protein levels are reduced in 231 cells treated with increasing concentrations of R1i (Lovastatin) as confirmed via western blot. GAPDH was probed as loading control. **D.** 231 cells overexpressing MNRR1 show increased capacity to form mammospheres. *Left,* representative image of mammospheres. Cells were imaged at 4X, and scale bar represents 100μM. *Middle,* graph depicting the number of mammospheres formed in MNRR1 overexpressing cells as compared to EV overexpressing cells (n=16, *p<0.05). *Right,* western blot confirming that markers of mammosphere formation ALDH1A1 and CD44 are higher in the MNRR1 overexpressing cells. GAPDH was probed as loading control.

The HMG-CoA reductase inhibitor Lovastatin was identified as a potent inhibitor of MNRR1 transcription as measured by an *MNRR1-*luciferase construct stably transfected into breast cancer cells from a 2400 compound library of small molecules and natural products (**Supplementary Fig. S1c**). Lovastatin reduced cell growth in 231 and 468 cells with an IC50 of 7.97 μM (**Fig. 2B**) and 19.6 μM (data not shown). Lovastatin treatment reduced MNRR1 protein level in 231 (**Fig. 2C**) and 468 cells (**Supplementary Fig. S1d**). Formation of mammospheres has been shown previously to be a simple *in vitro* approach to evaluate the tumor initiation capacity of cells [24] and MNRR1 was identified as one of the most upregulated proteins in breast cancer cell mammospheres [25]. To test if MNRR1 *induces* mammosphere formation, we transfected 231 cells with either an empty vector or an expression plasmid for wild-type MNRR1 and evaluated mammosphere formation as detailed in the materials and methods. We found that cells overexpressing MNRR1 show a significant increase in their ability to form mammospheres. These cells also display an increase in the levels of mammosphere markers such as CD44 and ALDH1A1 (**Fig. 2D**).

### MNRR1 induces and is required for the EMT phenotype in TNBC cells

The EMT program occupies the centerstage of cancer metastasis. It is an evolutionarily conserved program that allows cancer cells to lose their polarity and adhesive properties and acquire a mesenchymal phenotype to facilitate metastasis [26]. As shown in **Fig. 3A**, 10A cells overexpressing wild-type MNRR1 display a pro-EMT phenotype with increased levels of beta-catenin, vimentin, and slug while decreasing levels of zona occludens 1 (ZO-1). To further characterize the role of MNRR1 in the EMT induction, 10A cells were exposed to TGF-β to induce EMT *in vitro* [27, 28]. TGF-β treatment led to increased MNRR1 mRNA and protein (**Fig. 3B and 3C**). Interestingly, MNRR1 is upstream of the markers. As shown in **Fig. 3D**, cells treated with TGF-β display an increase in the protein levels of both MNRR1 and Slug whereas both increases are abrogated in the presence of Lovastatin. MNRR1 inhibition by Lovastatin treatment also reduces protein levels of Slug (**Fig. 3E**). Furthermore, in matrigel cell invasion assays, Lovastatin treatment significantly reduced the invasive capacity of 231 cells in a concentration dependent manner (**Fig. 3F**). Transient transfection of a transcriptionally defective (TD-R1) [22] MNRR1 construct reduced vimentin expression in 468 cells, further implicating a role for MNRR1 in EMT signaling (**Supplementary Fig. S1e**). Stable siRNA knockdown of MNRR1 in 468 cells further demonstrated a reduction in the levels of N-Cadherin (**Supplementary Fig. S1f**), suggestive of a defective EMT phenotype. In addition, 231 cells expressing TD-R1 MNRR1 had reduced levels of the EMT marker SNAIL and increased ZO-1 (**Supplementary Fig. S1g**). Taken together, these results suggest that MNRR1 is required for an optimal EMT in response to TGF-β.

**Figure 3:**
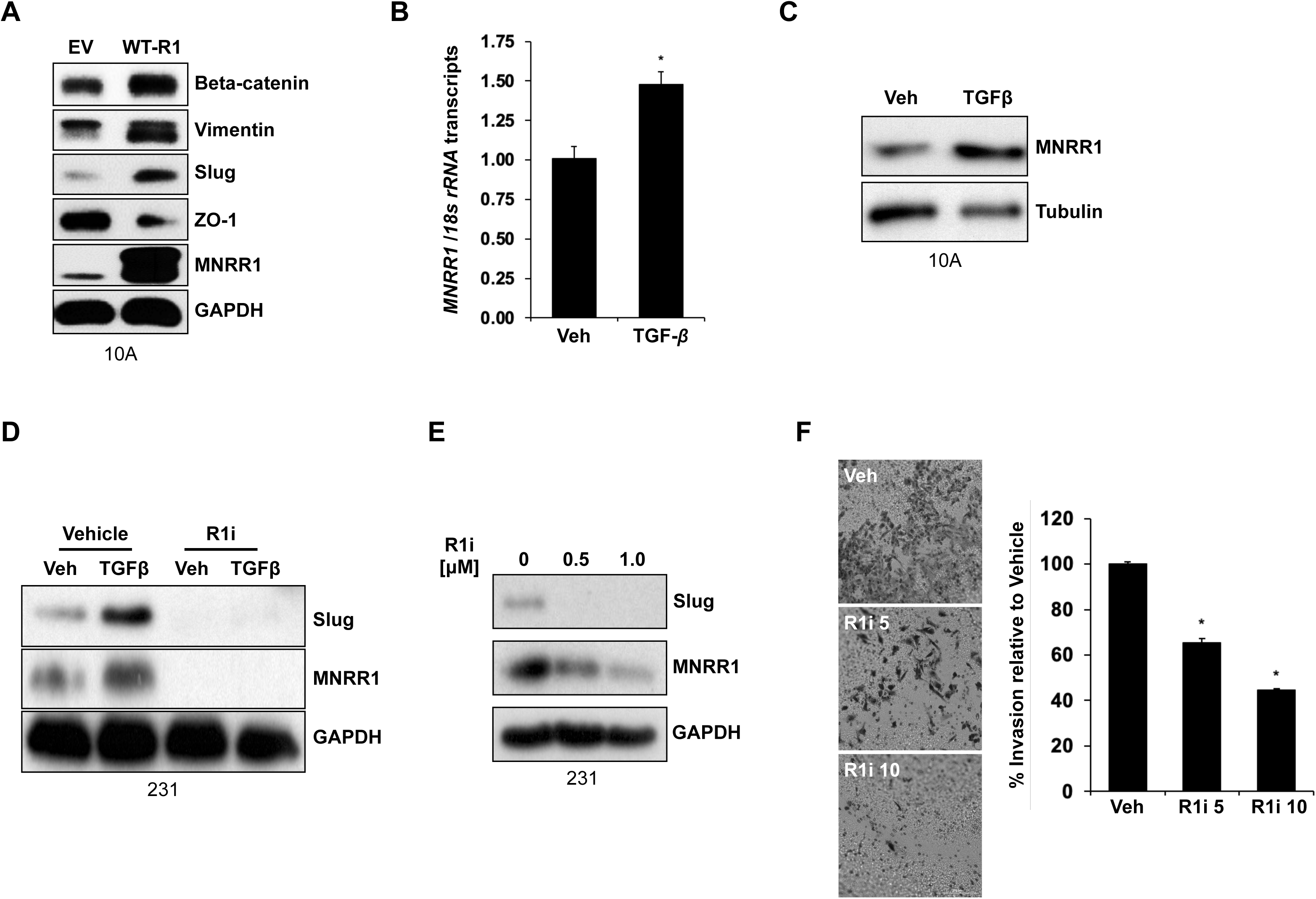
MNRR1 induces and is required for the EMT phenotype in TNBC cells. **A.** Western blot analysis of 10A cells overexpressing MNRR1 shows increased levels of mesenchymal markers (Beta-catenin, Vimentin, and Slug) as well as reduced levels of epithelial marker ZO-1 when compared to empty vector (EV) -overexpressing cells. These suggest an epithelial to mesenchymal transition (EMT). MNRR1 was probed to confirm overexpression and GAPDH was probed as loading control. **B.** Transforming growth factor beta (TGF-β) (5 ng/mL) increases the transcript levels of *MNRR1*. *18s rRNA* was used as a housekeeping gene for the RT-qPCR analysis (n=8, *p<0.05). **C.** TGF-β increases the protein levels of MNRR1 in 10A cells analyzed using western blot. Tubulin was probed as loading control. **D.** TGF-β increases the protein levels of mesenchymal marker Slug and the increase is blocked by co-treatment with MNRR1 inhibitor (lovastatin, 10 μM) as analyzed by western blot. MNRR1 was probed to confirm inhibition and GAPDH was probed as loading control. **E.** Protein levels of the mesenchymal marker Slug are reduced in 231 cells treated with increasing concentrations of the R1i, lovastatin. MNRR1 was probed to confirm inhibition and GAPDH was probed as loading control. **F.** *Left,* representative image depicting the reduced invasion capacity of 231 cells by treatment with increasing concentrations of lovastatin. Cells were imaged at 4x, and scale bar represents 100μm. *Right,* graph shows quantification of cell invasion in 231 cells treated with lovastatin. (n= 4, *p<0.05).

To further characterize the effects of MNRR1 loss on TNBC cell phenotypes, we generated a CRISPR/Cas9 MNRR1 knockout in 231 cells. Two clones, knockout 1 and 2 (KO-1 and KO-2), displayed a significant reduction in *MNRR1* transcript levels (**Fig. 4A**) and virtually no protein (**Fig. 4B**) compared to control cells (WT MNRR1, selected upon transfection with scrambled CRISPR plasmids). Loss of MNRR1 in KO-1 and KO-2 cells resulted in a significant reduction in mitochondrial function as well as reduced transcriptional activation of a luciferase reporter harboring an oxygen responsive element (ORE) that was previously shown to be activated by MNRR1 [29] (**Supplementary Fig. S2a**). We therefore used WT and KO-1 to characterize the effects of MNRR1 in 231 TNBC cells. KO-1 cells displayed significantly reduced cell growth (**Fig. 4C**), consistent with our previous observations, including on cell growth [12]. The doubling time of the WT cells was ∼27 hrs whereas that of KO-1 was ∼33 hr. Furthermore, KO-1 cells were significantly less invasive in matrigel assays (**Fig. 4D**). Slug is a vital EMT regulator that functions upstream of other EMT markers (*e.g.*, vimentin and cadherin) [30]. Slug is known to regulate EMT in multiple cancers such as lung, breast, ovarian, and pancreatic [31–34]. The transcript levels of *Slug* were reduced by ∼50% in the KO-1 cells (**Fig. 4E**). Slug protein levels were also found to be lower (**Fig. 4F**). Additionally, overexpressing wild-type Slug or wild-type MNRR1 using plasmids in KO-1 cells significantly enhanced the invasive capacity of the cells *in vitro* (**Supplementary Fig. S2b**). To identify the mitonuclear contribution of MNRR1 towards increase in Slug levels, we transfected either an empty vector, wild-type MNRR1, a transcriptionally defective mutant, or a transcriptionally superior mutant in KO-1 cells and evaluated for the levels of Slug. Only the wild-type and the transcriptionally active mutant of MNRR1 induce increased Slug levels, indicating that the effect of MNRR1 on Slug is via its nuclear function as a transcriptional regulator (**Supplementary Fig. S2c)**. Additionally, expressing the transcriptionally active version of MNRR1 in 231 cells significantly enhanced their invasive ability (**Supplementary Fig. S2d**). These results indicate that MNRR1 is upstream of Slug, and that MNRR1-mediated decreases in cell invasion occur at least in part via Slug depletion. To identify genes regulated by MNRR1, we performed RNA-Sequencing of WT and KO-1 cells. In the absence of MNRR1, 2857 genes were significantly downregulated (**Supplementary Fig. S3a)**. Scrutiny of the pool of downregulated genes using EnrichR revealed EMT to be one of the most significantly affected pathways (**Supplementary Fig. S3b**).

**Figure 4:**
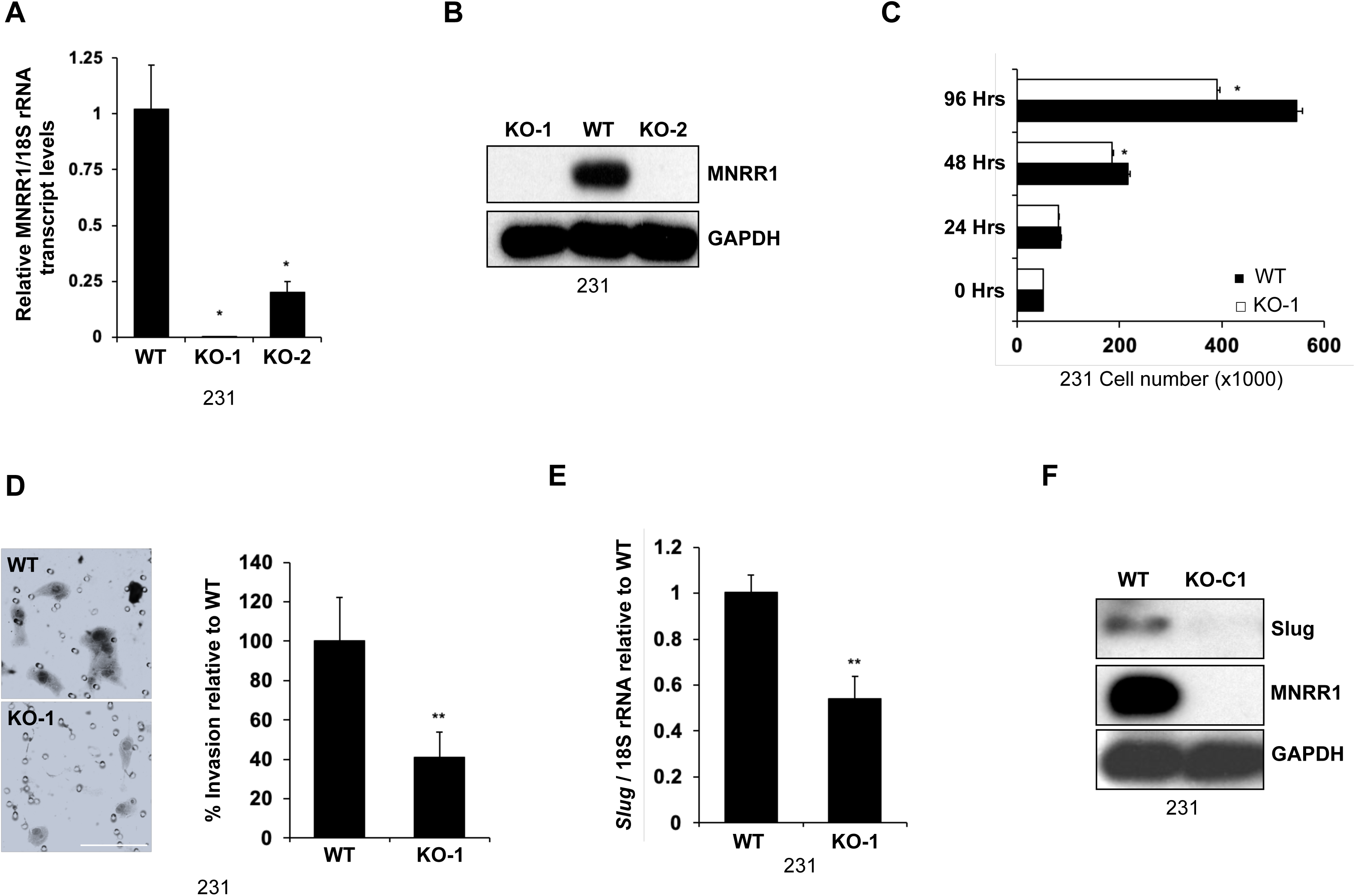
Loss of MNRR1 in TNBC reduces cell growth and invasion capacity. **A.** Transcript levels of *MNRR1* in 231 WT and two clones (KO-C1 and KO-C2) to confirm knockout. *18s rRNA* was used as a housekeeping gene for the RT-qPCR analysis (n=4, *p<0.05). **B.** Protein levels of MNRR1 in 231 WT, KO-C1, and KO-C2 to confirm knockout. GAPDH was probed as loading control. **C:** Cell growth is reduced in KO-C1 cells compared to WT, measured using cell number using trypan blue over a period of 96 h (n=4, *p<0.05). **D.** *Left,* Representative image depicting reduced cell invasion capacity of 231 KO-C1 cells. *Right,* graph showing quantification of cell invasion of 231 WT and KO-C1 knockout cells. Cells were imaged at 60x, and scale bar represents 50μm (n=8, **p<0.01). **E.** Transcript levels of *Slug* are reduced in KO-C1 cells. *18s rRNA* was used as a housekeeping gene for the RT-qPCR analysis (n=6, **p<0.01). **F.** Protein levels of Slug are reduced in KO-C1 cells. GAPDH was probed as loading control.

### Anti-apoptotic function of MNRR1 contributes towards the oncogenic phenotype *in TNBC*

MNRR1 is known to interact with Bcl-xL to inhibit Bax oligomerization and, subsequently, apoptosis [15], and we previously reported that phosphorylated MNRR1 in the mitochondria interacts with COX to activate OxPhos [21, 35]. We therefore hypothesized that the non-phosphorylated mitochondrial pool of MNRR1 preferentially interacts with Bcl-xL, localized on the outer mitochondrial membrane, to inhibit apoptosis. Daunorubicin is an anthracycline chemotherapeutic widely used in the treatment of TNBC [36]. Treatment of both WT and KO-1 231 cells with daunorubicin resulted in a concentration-dependent decrease in cell growth, with KO-1 displaying enhanced sensitivity (**Fig. 5A**), suggesting, MNRR1 expression may negatively regulate apoptosis. Daunorubicin treatment reduced the protein levels of MNRR1 and increased levels of cleaved PARP (cPARP), a marker of apoptosis (**Fig. 5B**). Furthermore, KO-1 cells displayed an increase in levels of cPARP compared to WT, indicating increased sensitivity to apoptosis (**Supplementary Fig. S3c**). Daunorubicin treatment also reduced MNRR1 levels in 468 cells, indicating loss of MNRR1 to be a phenotype of cells undergoing apoptosis (**Supplementary Fig. S3d**). To test our hypothesis that non-phosphorylated pool of MNRR1 interacts with Bcl-xL, 231 cells were transfected with 3xFlag-tagged expression plasmids encoding phosphomimetic (Y99E) MNRR1 or non-phosphorylatable (Y99F) MNRR1 and assessed for the ability to interact with co-transfected HA-tagged Bcl-xL. Y99F preferentially interacted with Bcl-xL (**Fig. 5C**) and displayed reduced cPARP levels when exposed to Daunorubicin (**Fig. 5D**). During intrinsic apoptosis, pro-apoptotic BCL2 family proteins (*i.e.*, Bax and Bak) oligomerize on the OMM and induce a phenomenon known as mitochondrial outer membrane permeability (MOMP) that facilitates the release of cytochrome *c* (CYCS) into the cytosol. Cytosolic cytochrome *c* leads to the formation of the apoptosome and, subsequently, the cleavage of caspase 3 and PARP prior to cell death (reviewed in [37]. The channel/s through which CYCS is released have been shown to be composed of Bax [38] and Bak [39], and also of the OMM protein VDAC [40–42]. Anti-apoptotic BCL2 family proteins (e.g. BCL2, MCL-1 and as Bcl-xL) function by sequestering pro-apoptotic proteins and preventing MOMP [43, 44]. The anti-apoptotic proteins also interact with VDAC which further limits the release of CYCS to prevent apoptosis [45, 46]. We, therefore, sought to identify whether non-phosphorylated MNRR1 is an integral component of the anti-apoptotic complex involving Bcl-xL and VDAC. Consistent with this notion, VDAC co-immunoprecipitated with Bcl-xL only in the presence of non-phosphorylated (Y99F) MNRR1 (**Fig. 5E**). These results indicate that non-phosphorylated MNRR1 plays a role in the anti-apoptotic pathways conferring resistance to apoptotic compounds.

**Figure 5:**
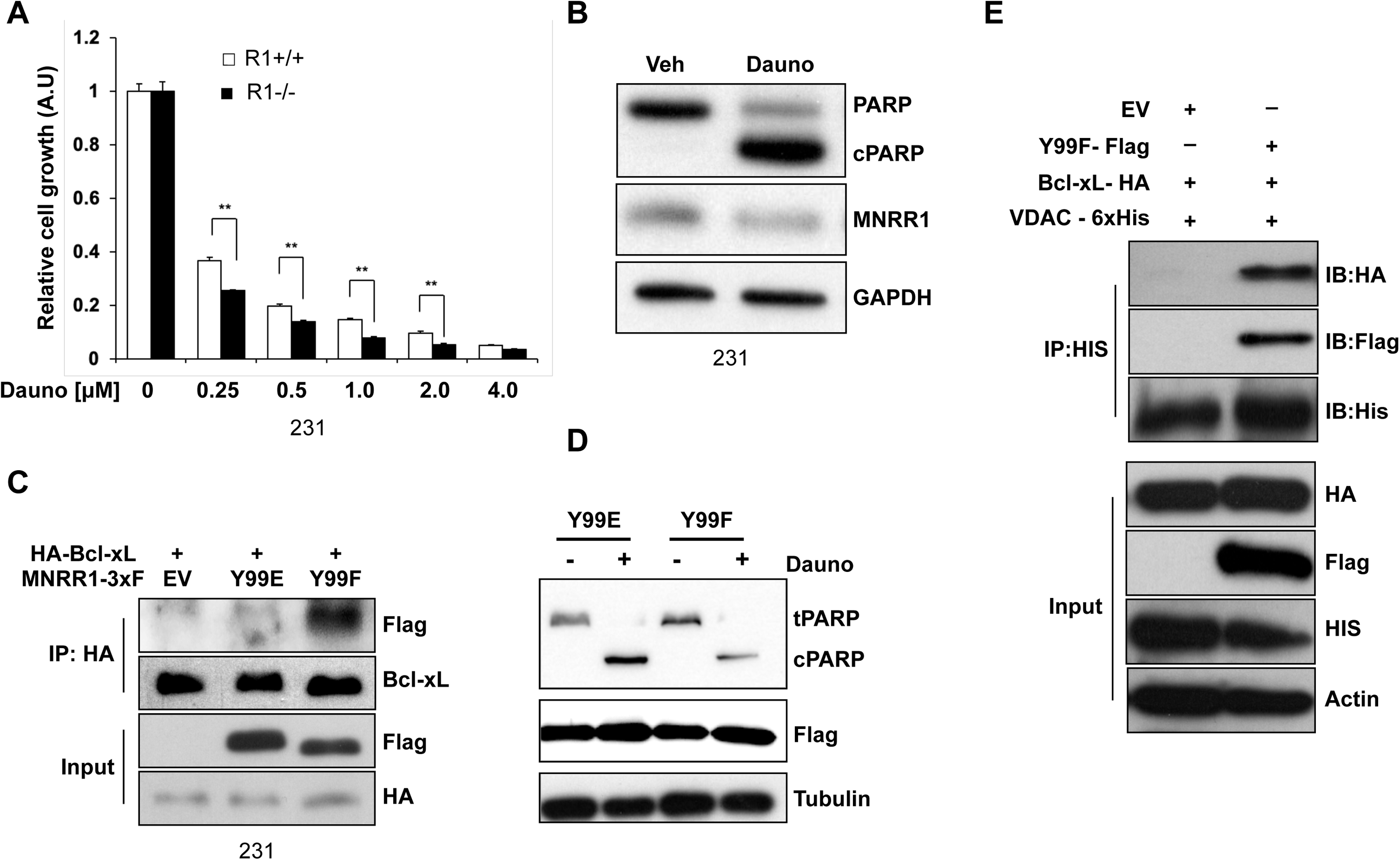
Anti-apoptotic function of MNRR1 contributes towards the oncogenic phenotype in TNBC. **A.** KO-C1 cells display defective cell growth relative to WT upon treatment with indicated concentrations of daunorubicin, measured using RealTime-Glo™ assay (n=8, **p<0.01). **B.** Western blot showing that MNRR1 protein levels are reduced during apoptosis induced by daunorubicin in 231 cells. PARP cleavage was assessed to confirm induction of apoptosis and GAPDH was probed as loading control. **C.** Co**-**immunoprecipitation (Co-IP) of HA-tagged Bcl-xL and either the FLAG-tagged phosphorylated (Y99E) or non-phosphorylatable (Y99F) MNRR1 mutants, showing that Y99F preferentially interacts with Bcl-xL. EV (empty vector) serves as a negative control. Input levels also shown. **D.** 231 cells overexpressing non-phosphorylatable (Y99F) MNRR1 are protected from daunorubicin-induced apoptosis compared to phosphorylated mimetic (Y99E) MNRR1. FLAG-tag MNRR1 was probed to confirm equal protein expression and Tubulin was probed as loading control. **E.** Co**-**immunoprecipitation showing that His-tagged VDAC1, HA-tagged Bcl-xL, and FLAG-tagged Y99F-MNRR1 form a complex. Empty Vector (EV) serves as a negative control. Equal amount of input lysates were probed to confirm expression of tagged proteins.

### Heat-Shock Factor 1 (HSF1) is upstream and drives the transcription of MNRR1 in TNBC

The aforementioned observations suggest that *MNRR1* is both transcriptionally and translationally increased in TNBC cell lines and is likely a significant contributor to the metastatic and resistance (to drug-induced cell death) phenotypes observed in TNBC patients. In addition, the MNRR1-promoter luciferase construct displayed enhanced activity in the HTB126 cancer cells compared to the HTB125 isogenic controls (**Fig. 1A**). To identify upstream signals that might transcriptionally regulate the *MNRR1* promoter, *MNRR1*-luciferase constructs were generated with deletions in five distinct regions (**Fig. 6A**) [47]. These mutants were tested in the isogenic breast-derived non-cancerous (HTB125) and cancerous (HTB126) cell lines. Promoter region deletion between 401-600 (Δ401-600) led to reduced *MNRR1*-luciferase activation in the HTB126 TNBC cell line. The other four deletions increased luciferase expression above the level observed in the HTB125 cells (**Fig. 6B**). These results suggested that the activating transcriptional signature for increased MNRR1 levels in TNBC is regulated at the 401-600-bp region on the *MNRR1* gene promoter. The Genomatix analysis Suite identified 55 *bonafide* transcription factor binding sites in the 401-600 region (not shown). Surprisingly, Heat-Shock Factor 1 (HSF1) was the only one in common between the Genomatix pool and previously published reports of proteins upregulated in 14 TNBC cell lines and four TNBC patient samples [48]. HSF1 is known to be a driver of transcripts that support malignant transformation [49]. To determine whether HSF1 could activate MNRR1, WT MNRR1-luciferase was co-transfected with an HSF1 expression plasmid or empty vector in MCF12A cells. HSF1 overexpression induced the WT-luciferase reporter ∼2-fold over that of empty vector expressing cells (**Fig. 6C)**. To extend and confirm this observation, enhanced HSF1 is also seen at the protein level in the transfected cells (**Fig. 6D).** We generated an *MNRR1* promoter luciferase construct with a mutation in the HSF1 binding site (HSF1mut) (**Fig. 6E**). HSF1 overexpression (HSF1-OE) increased reporter activity ∼15-fold and was blunted to almost half in the HSF1mut reporter plasmid (**Fig. 6F**). These results indicate that HSF1 is upstream of MNRR1 and drive its expression in TNBC cells.

**Figure 6:**
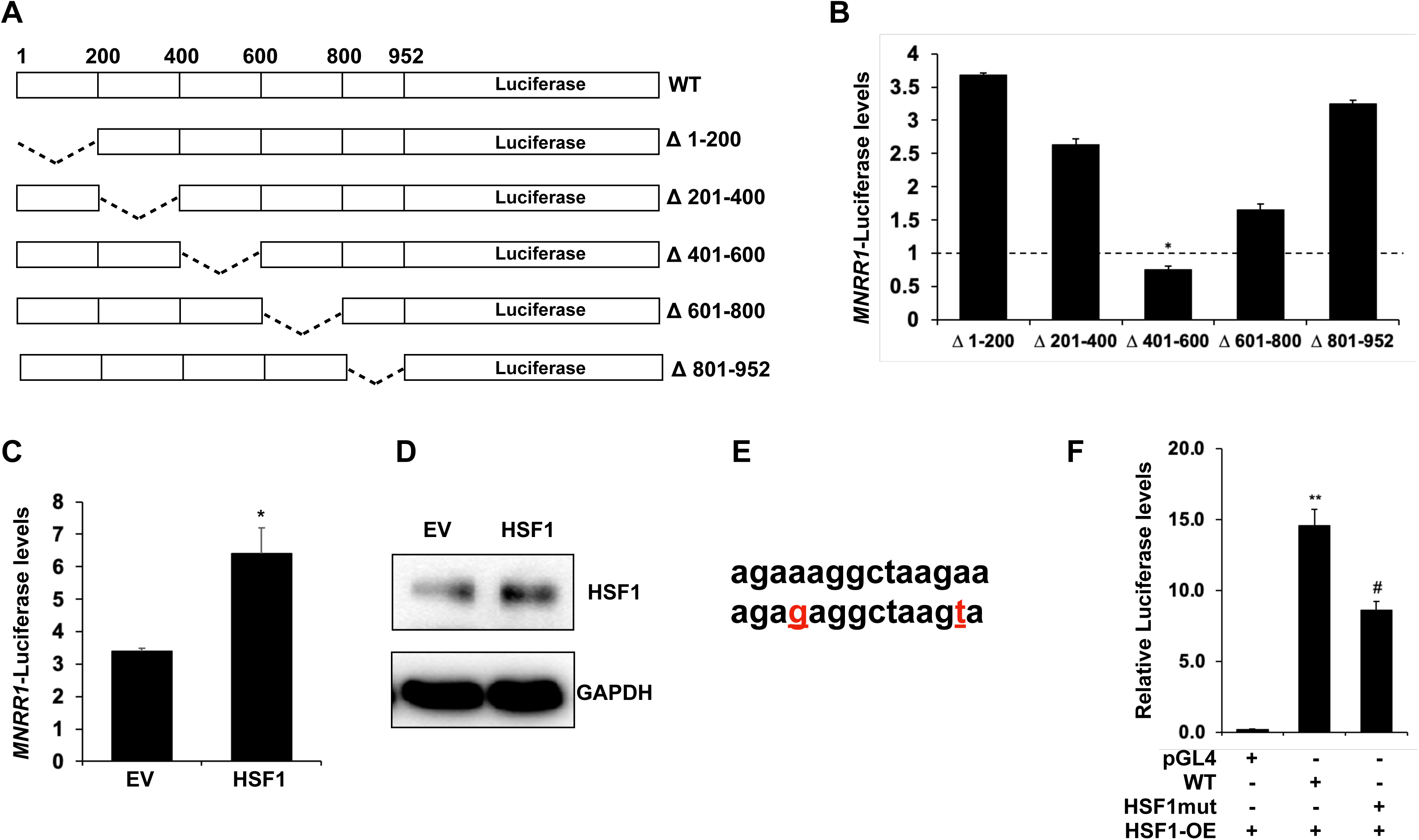
Heat-Shock Factor 1 (HSF1) is upstream and drives the transcription of MNRR1 in TNBC. **A.** Schematic representation of deleted regions of the *MNRR1* promoter. **B.** Dual luciferase reporter assay of various *MNRR1* promoter deletions showing that Δ 401-600 is not activated in HTB126 cells compared to HTB125 cells (dotted line), suggesting that the transcription factors that bind this region are required for activation of *MNRR1* in HTB-126 cells (n=4, *<0.05). **C.** Activation of full length *MNRR1*-promoter harboring luciferase reporter is higher in HSF1 overexpressing cells (n=4, *p<0.05). **D.** Western blot to confirm overexpression of HSF1 for C. GAPDH was probed as loading control. **E.** HSF1 binding site in the *MNRR1* promoter and the bases mutated (red, underlined) to generate the HSF1mut reporter plasmid. **F.** Mutant MNRR1 reporter (HSF1-mut) inhibits optimal activation of *MNRR1* using the dual luciferase reporter assay. pGL4 is the promoterless luciferase reporter that serves as a negative control. For all three groups n=8; ** p<0.01 between pGL4 and WT, # p<0.01 between WT and HSF1mut).

### MNRR1 protein levels are relatively higher in TNBC tissue samples and cell lines derived from Non-Hispanic Black patients

Our initial observation of higher MNRR1 transcript and protein levels in 468 cells compared to 231 cells (**Figs.1C and 1D**) led to the intriguing hypothesis that MNRR1 levels might be relatively higher in NHB women. The MDA-MB-468 and the MDA-MB-231 cell lines are representative of age-matched NHB and NHW TNBCs. We evaluated the levels of MNRR1 protein in de-identified TNBC tissues from NHB and NHW patients using immunofluorescence as detailed in the materials and methods. MNRR1 protein levels were significantly higher in the NHB patients (**Fig. 7A**). Analysis of *MNRR1* transcript levels in five non-cancerous breast tissue biopsy samples from de-identified NHB and NHW patients revealed ∼2-fold higher levels in NHB samples (**Fig. 7B)**. We then performed xenograft experiments using 468 cells and used another MNRR1 inhibitor, cephalexin, previously identified as reducing *MNRR1* promoter activity (**Supplementary Fig. S3e).** Finally, xenograft experiments were performed to analyze the effect of MNRR1 inhibition on tumor growth. Six mice were injected with 468 cells in a 1:1 matrigel mix. At the time of xenograft mice were either treated with 75 mg/kg oral cephalexin or vehicle by oral gavage twice a day. Tumor weight was evaluated at least every other day for 34 days. Cephalexin treated animals displayed an ∼50% reduction in tumor weight (**Fig. 7C**) and their excised tumor tissues contained reduced protein levels of MNRR1 (**Figs. 7D, 7E)**.

**Figure 7:**
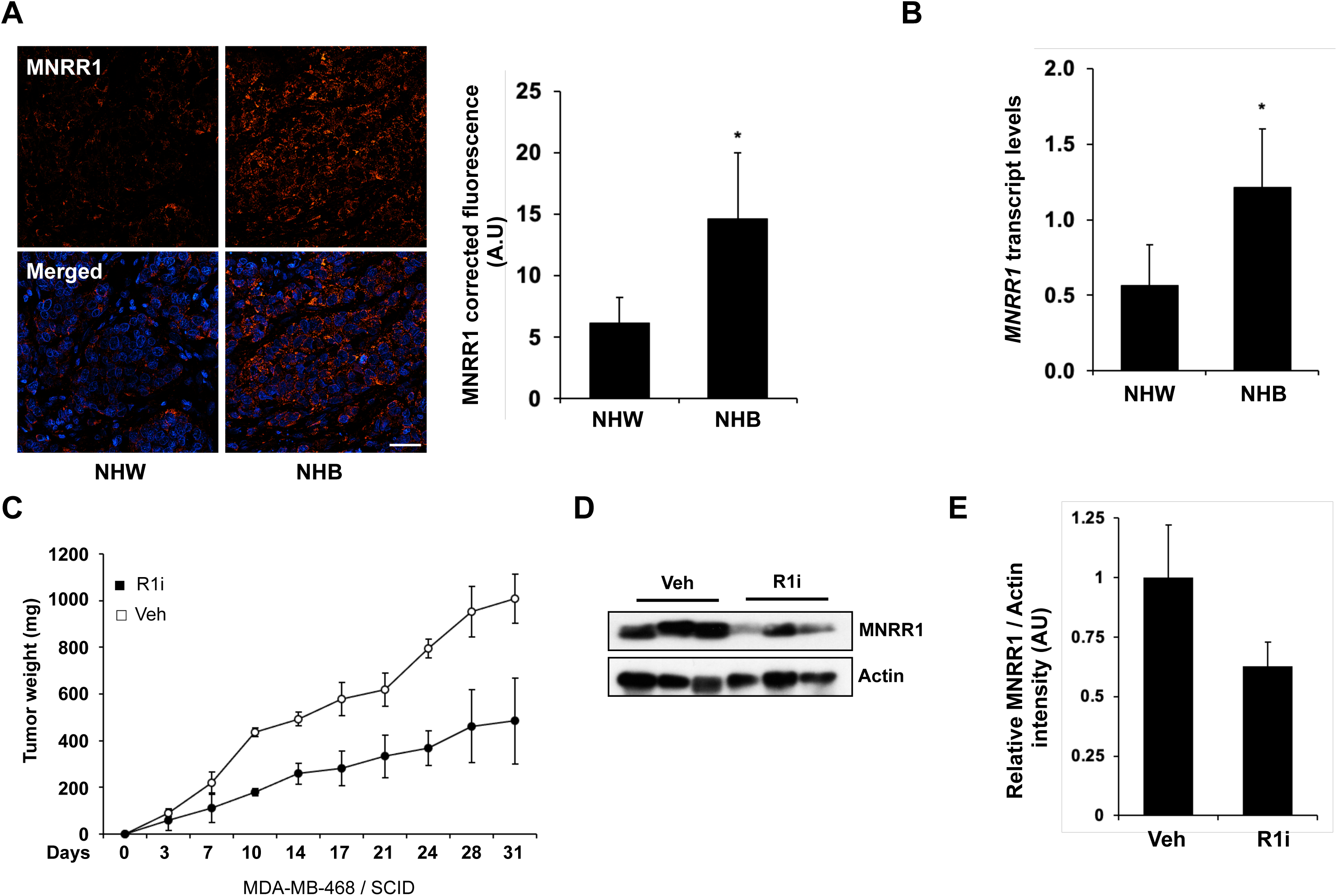
MNRR1 levels display a bias with higher levels in NHB relative to NHW patients. **A.** *Left*, Protein levels of MNRR1 are higher breast tissues sections from NHB compared to NHW detected via immunofluorescent staining. DAPI was used to label cellular nuclei. Tissues were imaged at 60x, and scale bar represents 20μm. *Left*, Quantification of protein levels from NHB and NHW tissues (n=14, *p<0.05). **B.** Transcript levels of *MNRR1* are higher in breast tissues from non-Hispanic Black women (NHB) compared to non-Hispanic White women (NHW). *18s rRNA* was used as a housekeeping gene for the RT-qPCR analysis (n=5, *p<0.05). **C:** Breast tumors grow more slowly in the presence of the MNRR1 inhibitor cephalexin as detailed in Materials and Methods (n=3 in each group). **D:** Protein levels of MNRR1 are reduced in tumors harvested from mice treated with vehicle or cephalexin. Actin was probed as loading control. **E:** Quantification of MNRR1 protein levels from harvested tumors in D.

## Discussion

MNRR1 is a bi-organellar regulator of cellular function. Although loss of function is associated with neurodegenerative phenotypes [50], increased levels and gain-of-function is associated with cancers [14, 16]. Triple-Negative Breast Cancer (TNBC) is a therapeutic challenge due to its frequent recurrence, the development of drug resistance, and enhanced aggressiveness; indeed, tumor metastasis is a principal factor driving mortality in breast cancer [51]. Chemotherapy and radiation, the mainstay approaches to TNBC, continue to provide unacceptably low response rates. Although mortality from breast cancer in general has declined by nearly 40% since 1990, certain subtypes, in particular TNBC, remain frustratingly high and disproportionately so for Non-Hispanic Black (NHB) women, where the incidence is roughly twice that of other ethnic groups [52]. Our data indicate that MNRR1 levels in TNBC cell lines and patient tumors are higher, specifically in tumor samples from NHB patients compared to NHW. MNRR1 is a pro-growth regulator and, consistent with our previously published work, MNRR1 levels are proportional to cellular growth profile both *in vitro* and *in vivo*. Here, using reporter deletion mutants, we have identified HSF1 as an upstream inducer of MNRR1 in TNBCs. HSF1 has been shown previously to regulate a transcriptional program in malignant cancers [53].

Work by us and others has identified three functions of MNRR1, two in the mitochondria and one in the nucleus. In the mitochondria, upon phosphorylation by Abl2 kinase on residue Y99, the interaction between phosphorylated protein and COX is enhanced, resulting in activation of the enzyme to increase ATP production [21]. The other mitochondrial function is the ability of MNRR1 to interact with Bcl-xL and inhibit apoptosis [15]. In the nucleus, MNRR1 interacts with RBPJκ to activate transcription [12, 29]. Deacetylation by Sirt1 makes the protein a superior transcriptional activator [22]. We recently uncovered that MNRR1 functions in the nucleus as a transcriptional co-activator by recruiting the histone acetyl transferase p300 to RBPJκ [54]. All these three roles contribute to a successful cancerous cell: (i) energy supply, (ii) prevention of apoptosis, thus allowing cells to grow in an uninhibited manner, and (iii) the transcriptional activation to induce a phenotype conducive for metastasis. We have here identified that it is the non-phosphorylated pool of MNRR1 in the mitochondria that preferentially interacts with Bcl-xL to prevent apoptosis. Non-phosphorylated MNRR1 is a part of an anti-apoptotic complex involving bonafide regulators such as Bcl-xL and VDAC so that loss of MNRR1, either by genetic manipulation or by using inhibitors such as lovastatin or cephalexin, sensitizes cells to apoptotic stimuli. This effect might be therapeutically exploited in drug resistant TNBCs. Furthermore, the nuclear function of MNRR1 as a transcriptional regulator contributes heavily to the EMT pathway. RNA-sequencing analysis identified this to be the most regulated pathway. *In vitro* analysis suggests that transcriptional regulation by MNRR1 is upstream of Slug, a key EMT pathway regulator. Additionally, loss of function models of MNRR1 display the inability of cells to invade through matrigel.

Finally, MNRR1 levels display a racial bias between TNBC tumor samples from NHB compared to NHW. Protein levels were found to be significantly higher in NHB tumors. Consistent with this observation, a SEER survey analyzing breast cancer cases from 39 states between 2010 – 2014 revealed that young NHB women have enhanced TNBC disease burden [52]. Attempts to describe the risk factors driving this racial disparity have been widely discussed in cellular and socio-economic terms but the outcome gap continues to be wide. Biologically it has been demonstrated that pro-growth signaling networks function aberrantly in NHB women with TNBC. Specifically: 1) BRCA1 loss of function mutations are higher [55]; 2) EZH2, a histone methyl transferase that functions by inhibiting BRCA1, is overexpressed [56]; and 3) anti-apoptotic AKT signaling is enhanced due to a loss of PTEN [57]. These studies suggest that, even in the absence of target receptors, there are molecular features of TNBC that might be therapeutically exploited. Taking into consideration the role of MNRR1 as an upstream inducer of the EMT pathway, disparity in the levels of MNRR1 between NHB and NHW groups could underlie aggressive disease and mortality observed in the former group [58]. Indeed, as shown in our data, MNRR1 levels are higher in apparently normal breast tissues from NHB individuals. Thus, higher levels of a pro-growth and anti-apoptotic gene could be responsible for the more aggressive disease observed in NHB population. Uncovering the upstream signal responsible for enhanced activation of MNRR1 in NHB populations is a clear need of subsequent research. The levels of MNRR1 across other racial groups are currently unknown, as is information about whether such levels co-relate with tumors in those groups. In summary, MNRR1 is an attractive therapeutic target in TNBCs both for primary as well as metastatic tumors. MNRR1 inhibitors identified among existing drugs clearly merit further study.

## Materials and Methods

### Cell Lines

The HTB125, HTB126, and MCF10A (10A) cells were purchased from ATCC (Manassas, VA, USA) and were grown in ATCC recommended media. The MDA-MB-231 (231) and MDA-MB-468 (468) cells lines were a kind gift from Dr. Kezhong Zhang (Wayne State University, Detroit USA) and were grown in DMEM with 1 mM pyruvate supplemented with 10% FBS (Sigma Aldrich, Burlington, MA, USA) plus penicillin-streptomycin. The MNRR1-KO (MDA-MB-231) cells were grown in DMEM with 1 mM pyruvate and 10% FBS (Sigma Aldrich, Burlington, MA, USA) plus penicillin-streptomycin and puromycin (1 μg/mL, Invivogen, San Diego, CA, USA).

### Chemicals

TGF-β was purchased from Invivogen (San Diego, CA, USA). Daunorubicin and lovastatin were purchased from MedChemExpress (Monmouth Junction, NJ, USA). Pharmaceutical grade cepahlexin was purchased from All Care Pharmacy, Allen Park, MI.

### Plasmids

The *MNRR1* 952-bp promoter luciferase reporter plasmid, pRL-SV40 *Renilla* luciferase expression plasmids, and the full length MNRR1-expressing plasmid have been described previously [47]. The promoterless luciferase reporter plasmid was obtained from Promega (Madison, WI, USA). The plasmids for expression of *MNRR1* promoter deletions (200 bp) driving the expression of lucifease were generated using overlap extenstion in WT-MNRR1 plasmid [47]. The transcriptionally active (TA, all lysine residues mutated to arginines), and the transcriptionally defective (TD, Δ101-113, deletion of amino acids 101-113) versions of MNRR1 have been previously described [12, 22]. The phosphomimetic (Y99E) and the non-phosphorylated mutants have also been previously described [21]. The pcDNA-hHSF1 WT was a gift from Lea Sistonen (Addgene plasmid # 71724) [59] and SlugMyc_pcDNA3 was a gift from Paul Wade (Addgene plasmid # 31698). The point mutations in the MNRR1 promoter HSF-1 site were generated using QuikChange Lightning Site-Directed Mutagenesis Kit (Agilent, Santa Clara, CA) and confirmed by sequencing. All expression plasmids were purified using the EndoFree plasmid purification kit from Qiagen (Germantown, MD, USA).

### Transient Transfection

Cells were transfected with the indicated plasmids using ViaFect or Transfast transfection reagent (Promega, Madison, WI, USA) according to the manufacturer’s protocol. A reagent–DNA ratio of at least 3:1 was used. Following incubation at room temperature for ∼15 min, the cells were overlaid with the mixture. The plates were incubated overnight at 37 °C followed by replacement with fresh complete medium and further incubation for the indicated time.

### Real-Time Polymerase Chain Reaction (RT-PCR)

Total cellular RNA was extracted from cells or tissues with a RNeasy Plus Mini Kit (Qiagen, Germantown, MD, USA) according to the manufacturer’s instructions. Complementary DNA (cDNA) was generated by reverse transcriptase polymerase chain reaction (PCR) using the ProtoScript^®^ II First Strand cDNA Synthesis Kit (New England Biolabs, Ipswich, MA, USA). Transcript levels were measured by real time PCR using SYBR green on an ABI 7500 system. Real-time analysis was performed by the ΔΔ^Ct^ method. The primers used were *MNRR1* forward: 5′-CACACATGGGTCACGCCATTACT-3′, reverse: 5′-TTCTGGGCACACTCCAGAAACTGT-3′; *Actin* forward: 5′-CATTAAGGAGAAGCTGTGCT-3′, reverse: 5′-GTTGAAGGTAGTTTCGTGGA-3′; *18s* forward: 5′-CCAGTAAGTGCGGGTCATAA-3′, reverse: 5′-GGCCTCACTAAACCATCCAA-3′.

### Luciferase Reporter Assay

Luciferase assays were performed with the dual-luciferase reporter assay kit (Promega, Madison, WI, USA). Briefly, cells were lysed in Promega 1x passive lysis buffer and 25 μL of lysate was used for assay with a tube luminometer using an integration time of 10 s. Transfection efficiency was normalized with the co-transfected pRL-SV40 Renilla luciferase expression plasmid [29].

### Cell growth

Cell growth was measured using RealTime-Glo™ MT Cell Viability (Promega, Madison, WI, USA). Briefly, 7.5×10^3^ cells were plated in a 96-well plate and treated with various concentrations of drugs for 0-72 h. Luminescence was measured using Promega GloMax Discover (Promega, Madison, WI, USA) with an integration time of 0.5 s. Cell counting was performed as described previously (and is reported as cell number x 1000 [12].

### Mammosphere formation assay

Mammosphere formation was performed in 231 cells using reagents and protocol from PromoCell (Heildelberg, Germany) per the manufacturer’s instructions. Spheroids were imaged at 4X or 20X using the BioTek Cytation C10 and the Gen5 software (version 3.11.19, Agilent, Santa Clara, CA) and counted manually.

### Cell invasion assay

Cell invasion was measured using Corning® BioCoat™ Matrigel® Invasion Chamber (Catalog #355480, Corning, NY, USA) per the manufacter’s instructions. Briefly, cell culture insert were rehydrated by placing them in a 24-well plate with complete medium for 2 hours in a humidified tissue culture incubator, 37 °C, 5% CO_2_ atmosphere. Post hydration, the wells were transferred to a plate with complete medium. 5×10^4^ cells (in serum free media) were added inside the inserts and they were incubated for 22 h in a humidified tissue culture incubator, 37 °C, 5% CO_2_ atmosphere. Cell culture inserts were removed, and the insides were scraped using cotton-tipped swabs moistened with medium. Next, the cells were fixed with methanol, stained with 1% toluidine blue, removed from the inserts using a scalpel, and mounted on glass slides with immersion oil. Cells were imaged at 4X, 20X, or 60X with the BioTek Cytation C10 as above and counted manually to calculate invasion compared to controls.

### Immunoblotting

Immunoblotting was performed as described previously [60]. Briefly, cell lysates for immunoblotting were prepared using RIPA buffer (Abcam, Waltham, MA, USA) and included a protease and phosphatase inhibitor cocktail (Sigma-Aldrich, St. Louis, MO, USA). Total protein extracts were obtained by centrifugation at 21,000 × g for 30 min at 4 °C. The clear supernatants were transferred to new tubes and quantified using the Bradford reagent with BSA as standard (BioRad, Hercules, CA, USA). Equal amounts of cell lysates were separated by sodium dodecyl sulfate–polyacrylamide gel electrophoresis (SDS–PAGE), transferred to PVDF membranes (BioRad), and blocked with 5% non-fat dry milk. Incubation with primary antibodies (used at a concentration of 1:500) was performed overnight at 4 °C. The ALDH1A1 (Catalog # 36671), PARP (9542), Beta-catenin (8480), Vimentin (5741), Slug (9585), ZO-1 (8193), Bcl-xL (24780), HA-tag (14031), HSF1 (12972), N-cadherin (13116), Snail (3789), GAPDH (8884), actin (12748), and tubulin (9099) antibodies were obtained from Cell Signaling (Danvers, MA, USA). The MNRR1 (19424-1) antibody was obtained from Proteintech (Chicago, IL). The FLAG antibody (A8592) was obtained from Sigma Aldrich (St. Louis, MO, USA) and the CD44s (MAB7045) was obtained from R&D Systems (Minneapolis, MN, USA). Incubation with secondary antibodies (1:5000) was performed for 2 h at room temperature. For detection after immunoblotting, the SuperSignal™ West Pico PLUS substrate or Super Signal™ West Femto Maximum Sensitivity Substrate (ThermoFisher, Waltham, MA, USA) was used to generate chemiluminescence signal, which was detected with X-Ray film (RadTech, Vassar, MI, USA) or BioRad Chemidoc digital imager (Hercules, CA, USA).

### Immunofluorescence

Cells plated on glass cover slips or breast tissue sections (Biobanking and Correlative Sciences Core, Karmanos Cancer Institute, Wayne State University) were fixed with 4% formaldehyde (prepared in 1x PBS) at room temperature for 15 min, followed by permeabilization with 0.15% Triton X-100 (prepared in distilled water) for 2 min, and then blocked with 5% bovine serum albumin (BSA) (prepared in 1x PBS, 0.1% TWEEN-20 (PBST) for 1 h at room temperature. Cells or tissue sections were washed with PBST then incubated for 1 h at room temperature in primary antibody solution containing Coralite® 594 conjugated mouse monoclonal anti-CHCHD2 IgG (1:100, Proteintech, Cat. No. CL594-66302) prepared in PBST. Cells or tissue sections were washed 3 times with PBST for 5 min each and mounted with Vectashield vibrance with DAPI (Cat. # H-1800-10, Vector Labs, Newark, CA, USA). Cells were imaged at 60x on the confocal 60 μm disk setting with the BioTek Cytation C10 as above). Six fields for each group were taken and z-stacks of 25 slices (±3 slices) were performed for each field followed by a z-projection and image deconvolution. Corrected total fluorescence for each field to determine MNRR1 content was calculated using FIJI (National Institutes of Health) (This work utilized the computational resources of the NIH HPC Biowulf cluster (https://hpc.nih.gov).

### Intact Cellular Oxygen Consumption

Cellular oxygen consumption was measured with a Seahorse XF^e^24 Bioanalyzer (Agilent, Santa Clara, CA, USA). Cells were plated at a concentration of 30,000 cells/well, a day prior to treatment and basal oxygen consumption was measured.

### Xenograft experiments

Six mice were injected with 2×10^6^ MDA-468 cells in 1:1 matrigel mix. At the time of xenograft mice were either treated with 75 mg/kg oral cephalexin or vehicle by oral gavage b.i,d. Tumor weight was evaluated at least every other day for 34 days. Tumors were harvested and analyzed for MNRR1 levels.

### RNA Sequencing and Analysis

Total cellular RNA was extracted using the RNEasy Plus kit (Qiagen, Germantown, MD, USA). Libraries with ribosomal depletion were prepared and sequenced on SP100 cycle flow cells. Raw paired-end RNA sequencing reads were aligned to the human reference genome (GENCODE v45 [61], primary assembly) with STAR [62]. Alignments with a mapping quality score below 10 were removed using SAMtools [63]. Gene-level quantification was carried out with the featureCounts function from the R/Bioconductor package Rsubread [64] using the GENCODE v45 gene annotation. Low-abundance genes with counts per million less than 1 in at least three samples were removed. Library size normalization was carried out with the TMM (Trimmed Mean of M-values) method [65] implemented in R/Bioconductor package, edgeR [66]. Differential expression analysis was performed in R using the limma–voom pipeline [67]. Linear models were fit to compare groups within each cell line, and significance was determined using moderated t test with Benjamini–Hochberg adjustment. Genes with adjusted p-value <0.1 and fold change >1.25 were considered differentially expressed. Genes with adjusted p-value < 0.1 and fold change > 1.25 were considered differentially expressed. Functional enrichment analysis of down-regulated genes was conducted with the clusterProfiler package [68] against MSigDB Hallmark gene sets.

### Statistical Analysis

All statistical analyses were performed with the two-sided Wilcoxon rank sum test using MSTAT version 6.1.1 (N. Drinkwater, University of Wisconsin–Madison).

## Supplementary Figure Legends

**Supplementary Figure 1.**
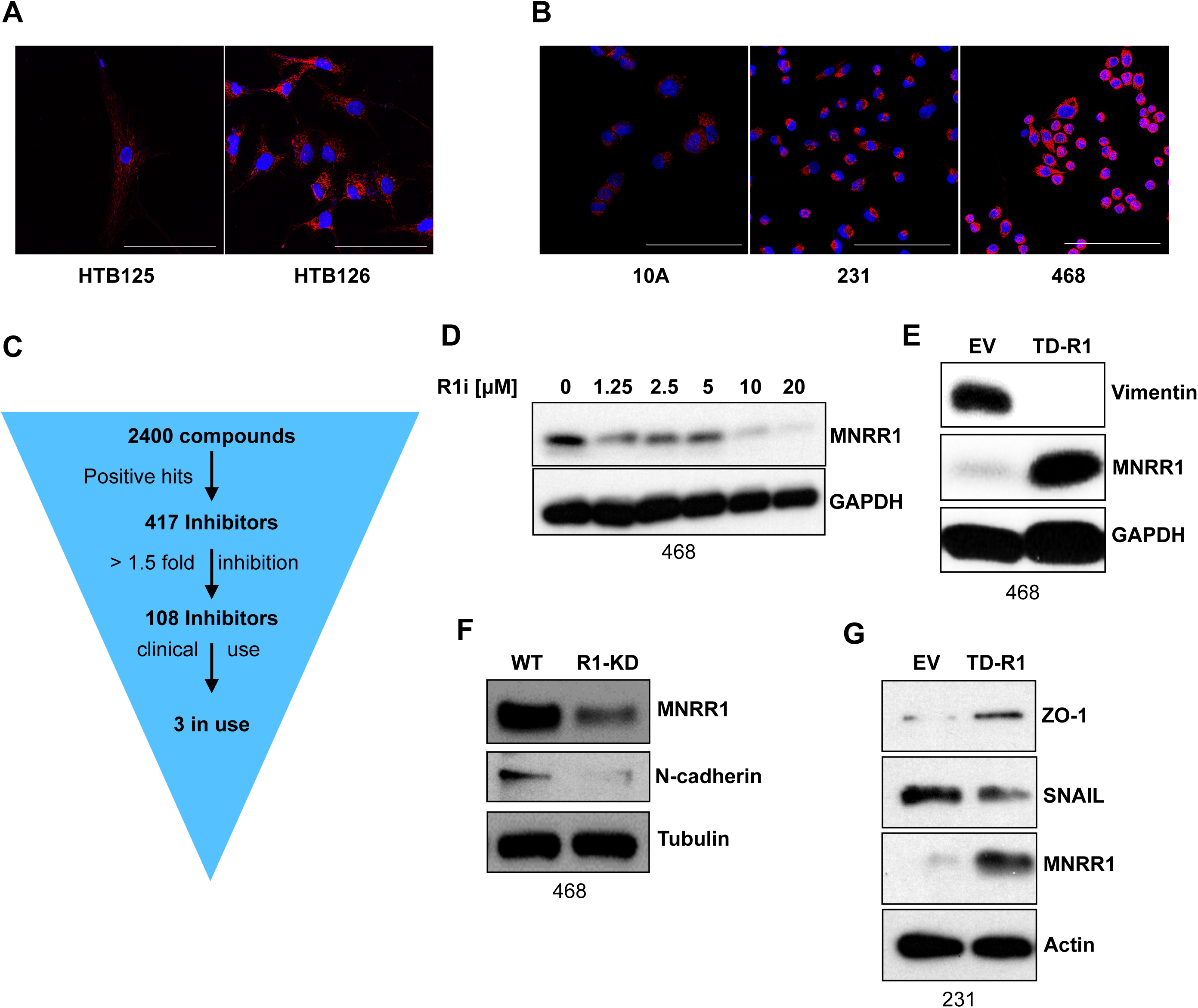
**S1A.** Protein levels of MNRR1 are higher in HTB126 (breast cancer line) vs HTB125 (isogenic control) cells detected via immunofluorescent staining. DAPI was used to indicate cellular nuclei. Cells were imaged at 60x, and scale bar represents 50μm. **S1B.** Protein levels of MNRR1 are higher in 231 and 468 (breast cancer lines) vs 10A (normal breast cell line) cells, detected via immunofluorescent staining. DAPI was used to indicate cellular nuclei. Cells were imaged at 60x, and scale bar represents 50μm. **S1C.** Overview of the high-throughput screening process to identify inhibitors of MNRR1 using the inverted pyramid approach. **S1D.** Protein levels of MNRR1 are reduced in 468 cells treated with increasing concentrations of MNRR1 inhibitor as analyzed via western blot. GAPDH was probed as loading control. **S1E.** Level of EMT marker vimentin is reduced in 468 cells overexpressing transcriptionally defective MNRR1 (TD-R1) compared to empty vector (EV). MNRR1 was probed to confirm overexpression and GAPDH was probed as loading control. **S1F.** Mesenchymal marker N-cadherin is reduced in MNRR1-knockdown (R1-KD) 468 cells as analyzed via western blot. MNRR1 was probed to confirm knockdown, and tubulin was probed as loading control. **S1G.** ZO-1 levels are increased, and Snail levels are reduced in 231 cells overexpressing transcriptionally defective MNRR1 (TD-R1) compared to empty vector (EV). MNRR1 was probed to confirm overexpression, and Actin was probed as loading control.

**Supplementary Figure 2.**
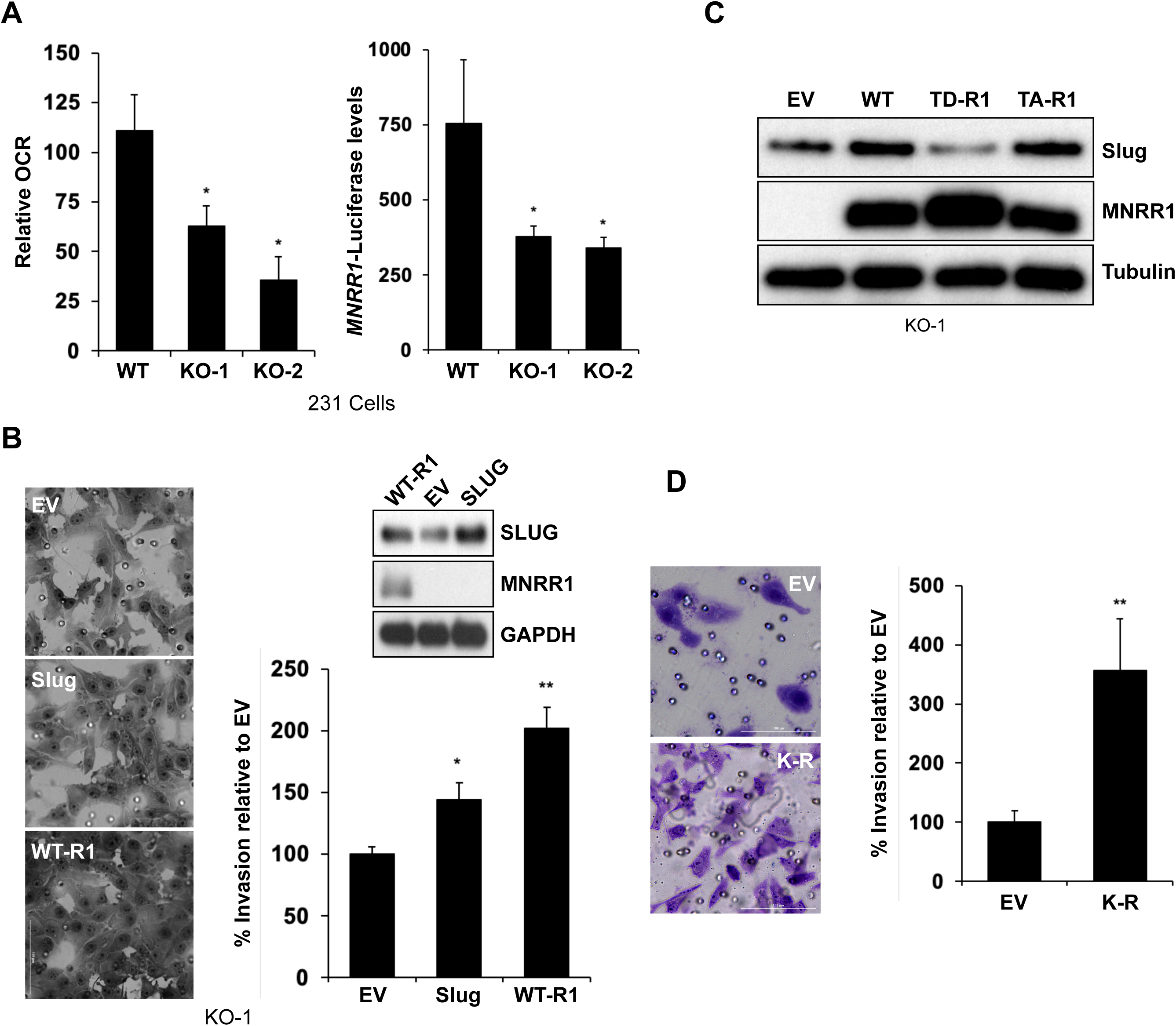
**S2A.** *Left,* Basal oxygen consumption in MNRR1 knockout clones KO-C1 and KO-C2 relative to WT (n=4, *p<0.05). *Right,* Activation of *MNRR1*-promoter luciferase reporterin KO-C1 and KO-C2 relative to WT, measured using dual luciferase reporter assay. (n=5, *p<0.05). **S2B.** *Left,* Representative image depicting increased cell invasion capacity of 231 MNRR1-knockout cells in Slug overexpressing cells and is even higher in MNRR1 overexpressing cells. *Right, above,* Western blot to confirm overexpression of MNRR1 and Slug. GAPDH was probed as loading control. *Right, below,* graph showing quantification of percent cell invasion relative to WT. Cells were imaged at 60x, and scale bar represents 100μm (n=5, *p<0.05, **p<0.01). **S2C.** Protein levels of Slug are modulated based on transcription function of MNRR1. MNRR1 knockout cells overexpressing WT (lane 2) and transcriptionally active MNRR1 (TA-R1, lane 4) have higher protein levels than EV (lane 1) and transcriptionally defective MNRR1 (TD-R1, lane 3). MNRR1 was probed to confirm overexpression and GAPDH was probed as loading control. **S2D:** *Left,* Representative image depicting that cell invasion capacity of 231 cells overexpressing transcriptionally superior MNRR1 (K-R MNRR1) is higher than EV expressing cells. *Right,* graph showing quantification of cell invasion. Cells were imaged at 60x, and scale bar represents 100μm (n= 5, **p<0.01).

**Supplementary Figure 3.**
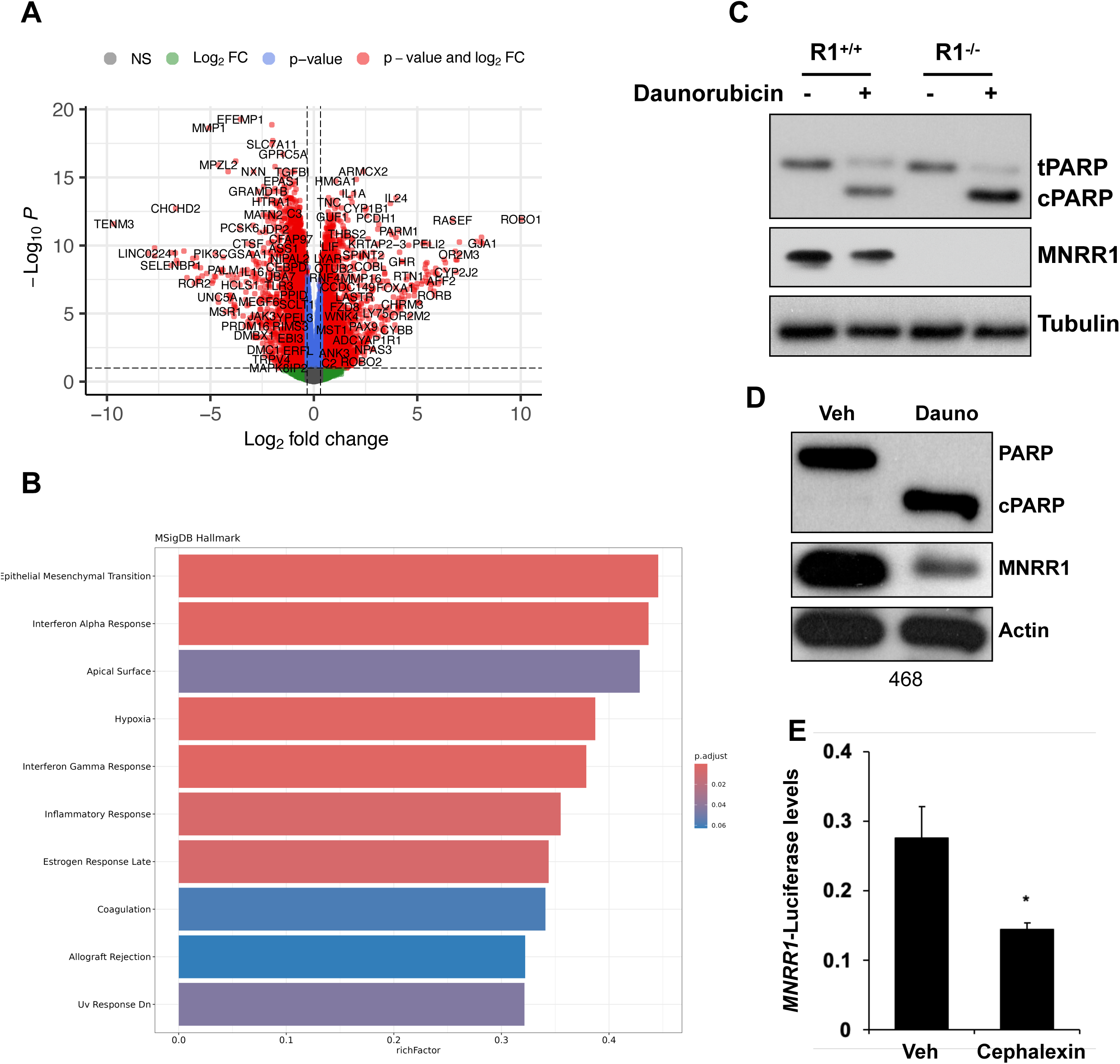
**S3A.** Volcano plot depicting differentially expressed genes in KO-1 cells compared to WT. **S3B:** Enrichment analysis of MSigDB Hallmark gene sets among downregulated genes identified in panel A. **S3C.** PARP cleavage is higher in 231 MNRR1-knockout cells treated with daunorubicin compared to WT. MNRR1 was probed to confirm knockout and Tubulin was probed as loading control. **S3D.** MNRR1 levels are reduced with daunorubicin treatment compared to vehicle (DMSO) in 468 cells. PARP cleavage was assessed to confirm induction of apoptosis and Actin was probed as loading control. **S3E.** Relative inhibition of *MNRR1*-promoter luciferase reporter in cells treated with either vehicle (DMSO) or 10μM of the MNRR1 inhibitor, cephalexin used in xenograft studies (n=4, *p<0.05).

## Acknowledgements

We would like to thank Dr. Julie Boerner, Director, Biobanking and Correlative Sciences Core, Karmanos Cancer Institute, Wayne State University, for deidentified TNBC patient samples. We would also like to thank Dr. Lisa Polin, Director, Animal model and therapeutics evaluation core, Karmanos Cancer Institute, Wayne State University for xenograft experiments.

## Author Contributions

M.S, P.M, C.V, M.MD, V.P, Y.X, S.R, M.P, H.E, S.Rai, S.R, N.P, performed experiments. G.B, A.T, N.P, S.A, analyzed data. N.P and S.A participated in experimental design. L.G and A.F provided intellectual feedback. N.P, A.F, L.G, S.A, wrote the manuscript. All authors have reviewed the manuscript.

## Data Availability Statement

All study data presented in this manuscript are included in the article or are available from the lead contact upon reasonable request. Unique reagents generated from this study are available from the lead contact with a completed Materials Transfer Agreement. This study did not generate original code.

